# Microplastics reach the brain and interfere with honey bee cognition

**DOI:** 10.1101/2023.08.05.552092

**Authors:** Elisa Pasquini, Federico Ferrante, Leonardo Passaponti, Francesco Saverio Pavone, Irene Costantini, David Baracchi

## Abstract

Scientific research exploring the impact of microplastics (MPs) in terrestrial systems is still at an early stage but has already confirmed that exposure to plastics leads to various detrimental health effects in several organisms. Although recent studies have shown the toxicological effects of single MP polymers on honey bees, the effects of different polymer combinations and their consequences on cognitive and behavioural performance remain unknown. To fill this knowledge gap, we investigated the effects of MPs, both individually and in combination, on the cognitive abilities of the honey bee *Apis mellifera*. We evaluated the acute oral toxicity of Polystyrene (PS) and Plexiglass (PMMA) MPs, as well as a combination of the two (MIX), at three different concentrations (0.5, 5 and 50 mg/L^-1^) and analysed their effects on sucrose responsiveness and appetitive olfactory learning and memory. We also explored whether these MPs could reach and accumulate in the insect brain using Two-Photon Fluorescence Microscopy (TPFM) in combination with an optimized version of the DISCO clearing technique. The results revealed that PS reduced the responsiveness of foragers to sucrose, whereas PMMA had no significant impact; however, the combination of PMMA and PS had a pronounced negative effect on sucrose responsiveness. In addition, both PMMA and PS, as well as MIX, impaired bee learning formation and memory retrieval, with PS exhibiting the most severe effects. Regarding our brain imaging analysis performed with TFPM, we found that after only three days of oral exposure, MPs could penetrate and accumulate in the brain. These results raise concerns about the potential mechanical, cellular, and biochemical damage that MPs may cause to the central nervous system.

## Introduction

In the not-too-distant future, part of the Anthropocene era will be likely be referred the “Age of Plastic”. Plastic products are widely used due to their practicality and convenience, making them the most ubiquitous materials in contemporary history (Cole et al., 2011; Millican & Agarwal, 2021). Despite the high benefits that our society has extolled, the conspicuous use of plastic has caused serious environmental problems, which have been exacerbated by the massive production of plastic objects (including primary microplastics) and inadequate disposal management (Cook & Halden, 2020). The primary issue with plastics is their poor biodegradability, which can take decades or even centuries to decompose (Weinstein et al., 2016), leading to the accumulation in the environment of extremely persistent particles known as microplastics (Ziani et al., 2023).

Microplastics (MPs) are polymeric particles of various shapes and compositions that are smaller than 5 mm and have been detected in nearly every environment on Earth (Rochman, 2018; Tursi et al., 2022; C. Wang et al., 2022). Although researchers are only beginning to assess the effects of MPs on terrestrial systems, studies suggest that exposure to plastic can have various negative consequences for organisms. These include upsetting the gut health balance (Huang et al., 2021; Lu et al., 2018), disrupting reproduction (Ilechukwu et al., 2022), altering behaviour (Chae & An, 2020), causing immunotoxicity, and disrupting gene expression (Muhammad et al., 2021; Sun et al., 2021). Once ingested, MPs can persist in the digestive systems of living organisms and accumulate at significant levels (Deng et al., 2021). Current research on the mosquito *Culex pipiens* has shown that MPs in water can be transferred from the early developmental stages to the pupal and adult stages of the insect, indicating the ability of MPs to move between different developmental stages (Al-Jaibachi et al., 2018, 2019).

The potential toxicity of MPs to insects is significant because these organisms are widespread in many environments and provide essential ecosystem services. In particular, pollinators such as bees have received special attention because of their steady decline. Recent studies have demonstrated the presence of MPs in honey (Diaz-Basantes et al., 2020; Liebezeit & Liebezeit, 2013) and in the cuticle of honey bees (Deng et al., 2021; Edo et al., 2021). These results are consistent with other studies suggesting that *Apis mellifera* can assimilate MPs from the environment through the cuticle or by ingestion (Alma et al., 2023; Buteler et al., 2022) and transfer them to other parts of the hive, such as the larvae, wax, and honey (Alma et al., 2023). These findings raise concerns because, besides the fact that bees can contaminate the human food chain through their products, exposure to these particles can have long-term negative effects on bee health and their ability to pollinate. To date, few studies have examined the potential risks posed by a limited types of MPs to bees. Most importantly, existing research only investigated the toxicological effects of single polymers, while animals, like honey bees are often exposed to multiple types of polymers simultaneously (Edo et al., 2021). MPs, especially those in polystyrene (PS) and polyethylene (PE), have been shown to adversely affect the survival of *A. mellifera* (Balzani et al., 2022; Deng et al., 2021; K. Wang et al., 2022), its immune response (K. Wang et al., 2022), the integrity of its gut microbiome (Deng et al., 2021; K. Wang et al., 2021, 2022), body weight (Al Naggar et al., 2023), and feeding behaviour (Al Naggar et al., 2023; Buteler et al., 2022). Furthermore, evidence shows that these particles accumulate in the haemolymph (Buteler et al., 2022; Deng et al., 2021) and likely spread and accumulate in various organs, including the brain. However, the impact of MPs on the behaviour and cognitive abilities of bees remains unclear despite the ongoing debate on their effects on bee health. So far, only one study has investigated the effects of acute and chronic exposure to different PE-MPs concentrations on bee cognition (Balzani et al. 2022). This study found only marginal effects on individual bees’ ability to respond consistently to sucrose, imposed only by the highest and ecologically irrelevant concentration tested (i.e., 50 mg L^-1^). However, even for this high concentration, no effects on bee learning or memory abilities were observed. Although Balzani et al. (2022) reported no cognitive impairments in the presence of environmentally relevant concentration of PE, it is possible that other micro polymers may impact bees’ health, brain and cognition. This is because, besides PE, pollinators commonly encounter many other polymers in the environment, such as Polypropylene (PP), Polyvinyl Chloride (PVC), Polystyrene (PS), Polyethylene Terephthalate, (PET), Polyurethane (PU), Acrylonitrile Butadiene Styrene (ABS), Polycarbonate (PC) and Polymethyl methacrylate (PMMA) among others (Edo et al., 2021). In addition, MPs that individually have negligible or minimal effects on animal health or cognition, PE included, may have amplified adverse effects when mixed with other MPs due to synergistic or additive interactions. Unfortunately, to our knowledge, interactions between different polymers have not been studied in any organism. The existence of this knowledge gap takes on even greater significance when considering the potential effects of these molecules on the vital cognitive processes that bees rely on, including learning, memory, and spatial navigation. These cognitive functions and flexibility are highly dependent on the integrity of neural structures and processes in the bee brain (Devaud et al., 2015; Giurfa, 2022; Menzel & Giurfa, 2006). Consequently, even minor impairments in cognitive function can profoundly affect the ability of individual bees to adapt and exhibit behavioural flexibility, thereby posing a significant threat to the survival of the entire colony (Klein et al., 2017). Similarly, they could disrupt the intricate network of interactions within the colony, leading to reduced cooperation, compromised division of labour, and reduced overall efficiency. Deficits can impact individual bees’ ability to adapt and demonstrate behavioural flexibility, ultimately putting the survival of the entire colony at risk (Baracchi, 2019). Given the severity of these potential consequences, our study aimed to enhance the understanding of the interactions between different MP polymers and their sublethal effects on bee cognitive processes. To this aim we conducted an acute oral toxicity test of Polystyrene (PS) and Polymethyl methacrylate/Plexiglass (PMMA) MPs, as well as a combination of the two, on individual bees to investigate any potential synergistic or additive effects. Additionally, we optimized a clearing procedure based on the DISCO technique to enable 3D visualization of intact bee brains at submicron resolution with TPFM (Renier et al., 2014). Our objectives were twofold: (1) to evaluate whether ingested MPs particles interfere with sucrose responsiveness and appetitive learning and memory, using established standard behavioural assays based on the proboscis extension response (PER), and (2) to investigate whether these MPs can reach and accumulate in the insect brain.

## Material and Methods

### Honey bees

All the experiments, both behavioural and neuronal, were carried out under laboratory conditions on the species *Apis mellifera ligustica*. Bees were collected from the experimental apiary, consisting of 6 colonies, and placed in front of the CBE Laboratory of the Department of Biology, University of Florence (Italy).

### Behavioural experiments

#### MPs solutions

Different groups of bees were exposed to PS (polystyrene), PMMA (Polymethyl methacrylate/plexiglass) or to a combination of both (hereafter ‘MIX)’ with the aim of investigating any potential additive or synergistic effects. Experimental MP solutions were prepared using PS (PSMS-1.07, 4.8-5.8 μm, Cospheric LLC, USA) and PMMA (PMPMS-1.2, 1-40 μm, Cospheric LLC, USA) nanospheres at three different concentrations (50, 5 and 0.5 mg L^-1^) in purified water with 50% (w/w) sucrose. These concentrations were designated as high (HC), medium (MC), or low (LC), and were chosen to represent the range of variability found in the natural environment (MC and LC), or a higher concentration (HC). These concentrations were also based on the only other published experimental study on the effect of MPs on honey bee cognition (Balzani et al., 2022), enabling a direct comparison of results. The sucrose solution used for the control bees did not contain MPs and was 50% (w/w) sucrose.

### MPs exposure and bees harnessing

All the behavioural tests were carried out between June and November 2022. To perform the experiments, foraging bees were captured at a feeder placed in front of the experimental beehive, providing 50 % sucrose solution (w/w), and randomly placed in bee cages (9 × 7 × 11 cm). The cages were immediately brought to the laboratory and equipped with 10 ml syringes that delivered the control sucrose solution (control group) or the MPs solutions (experimental groups) and were maintained in the dark at room temperature (23 ± 2 ◦C) and humidity (50 ± 10%) for 1 day. The next day, the bees were briefly chilled in ice for 5 minutes and individually harnessed in small cylinder holders by placing a tape strip in between the head and the thorax. To enable the free movement of their antennae and mouthparts and to observe their proboscis extension response (PER) to sucrose stimulation on the antennae, their heads were immobilised with low temperature melting wax. After recovery, all bees were administered 20 μl of control or experimental solution containing MPs, depending on their group, and kept overnight in a dark and humid environment at room temperature. On the following day, after a total of 48 hours exposure to the solutions, the bees were tested according to the experimental conditions (see below). For each of the MPs tested (PS, PMMA and MIX) in the two behavioural experiments, we had 1 control group and 3 experimental groups with different levels of MP exposure (LC, MC, HC).

### Sucrose responsiveness assay

The responsiveness of individual bees to sucrose is closely tied to their role in hive and foraging behaviours (Pankiw & Page Jr, 2000), and can be a good indicator of their cognitive performance when sucrose is used as a reward (Scheiner et al., 2001). The sucrose responsiveness of harnessed bees in relation to exposure to MPs was analysed by recording PER in response to increasing concentrations of sucrose, following (Baracchi et al., 2017). First, on the day of the test, we fed the harnessed bees early in the morning with 2μl of each respective solution. After 1 hour, prior to the test, bees were desensitized to water by touching the antennae with a soaked toothpick and allowed to drink *ad libitum*. Each bee was then presented with six sucrose solutions of increasing concentration in a logarithmic scale: 0.1%, 0.3%, 1%, 3%, 10%, 30%, with an inter-stimulus interval (ISI) of 2 minutes. To account for potential laterality in sucrose responsiveness, both antennae were stimulated using a wooden toothpick during each presentation (Baracchi et al., 2018). Sucrose presentations were interspersed with water stimulation to avoid sucrose sensitisation. Bees that did not react to any concentration of sucrose during the six stimulations were given a 50% sucrose solution at the end of the series. Bees that responded to the water stimulation or did not respond to the highest sucrose solution (50%), were not included in the analysis (Baracchi et al., 2017; Baracchi et al., 2018). The number of PERs shown in the 6 sucrose stimulations was used to calculate the individual sucrose response score (SRS). Replicates were performed for PS, PMMA, and MIX particles. A total of 374 bees were tested to evaluate the effects with PS (C, n = 85, LC: n = 100, MC: n = 103, HC: n = 86; a total of 348 bees belonging to the PMMA treatment (C: n = 83, LC: n = 94, MC: n = 92, HC: n = 79) and a total of 286 bees to MIX (C: n = 64, LC: n = 69, MC: n = 52, HC: n = 56,).

### Associative olfactory learning and memory recall test

The proboscis extension response (PER) is a reflexive feeding behaviour that is elicited when the bee’s antennae are stimulated with sucrose. We used a type of conditioning known as “differential conditioning” of the PER to assess the impact of MPs on bee learning and memory. This method involved exposing the bees to five combinations of a specific odorant (conditioned stimulus, CS+, either 1-hexanol or nonanal) paired with a 30 % sucrose solution (unconditioned stimulus, US) and to five presentations of another odorant without sucrose (CS-, either nonanal or 1-hexanol) (Baracchi et al., 2020). Consequently, the bees had to learn to react to the first odour (CS+) but not respond to the second (CS-). After 48 hours of exposure to the solutions (MPs and control), on the day of the experiment, the harnessed bees were fed early in the morning with 2μl of each experimental and control solution. After 1 hour, they were subjected to the conditioning test. Bees that showed a PER in response to the first presentation of CS+ were immediately excluded from the study, as learning could not have been assessed. Our experimental design was balanced by exposing half of the bees in each group to 1-hexanol as the reinforced odorant (CS+), while the other half received nonanal. To deliver these two odours to the bees, we used an odorant dispenser controlled by an Arduino Uno microcontroller. Training was initiated by placing a single bee 2 cm from a device which emitted a constant stream of clean air. After a period of 10 seconds to familiarise with the environment, a conditioned stimulus (either 1-octanol or nonanal) was presented for 4 seconds. During the reinforcement phase, 3 seconds after the onset of odour presentation, the bee’s antennae were stimulated with sucrose using a toothpick and allowed to drink for 3 seconds. After 10 seconds of rest, the bee was removed from the apparatus and replaced with another subject for processing. During the 10 conditioning trials, the occurrence of PER was recorded each time the conditioned stimuli (CSs) were presented to each individual. After the conditioning phase, the bees were placed in darkness and subjected to a memory test after 2 hours (middle-term memory, MTM) and 24 hours (early long-term memory, eLTM), in which odorants were provided without any reward. Additionally, the presentation order of CS+ and CS-was randomized among the bees. To evaluate memory retention, the percentage of bees in each group that responded correctly to CS+ and not CS- - the bees with specific memory - was quantified. Bees that did not respond during the memory retention test were given additional antennal stimulation with 50% sucrose solution to verify their motivation or physical condition. As specified in the previous behavioural experiment, replications were conducted for each type of MPs particles. In total, a number of 290 tested bees were exposed to PS (LC: n = 67, MC: n = 92, HC: n = 68, Control: n = 63), a number of 301 bees were exposed to PMMA (LC: n = 71, MC: n = 88, HC: n = 69, Control: n = 73), and a number of 347 bees were treated with MIX (LC: n = 73, MC: n = 111, HC: n = 83, Control: n = 80).

### Brain imaging experiments

#### Exposure to fluorescence MPs

Foragers were caught at a feeder and immediately randomly housed in bee cages (see above for details) equipped with 10 ml syringes containing either plain sucrose solution (control bees) or sucrose solution laced with red fluorescent polymer microspheres (FMR, 1-5 μm, Cospheric LLC, USA, 50 mg L^-1^ (exposed bees)). Exposure lasted 3 days. A total of 6 bees were treated to evaluate the presence of MPs in the brain (3 controls and 3 FMR administered bees).

### Brain preparation and iDISCO clearing protocol

Bees were sacrificed by CO_2_ flux and whole brains were dissected from the head capsule in PBS under a Leica Stereozoom S9i. Ocelli and pigmented cells of the optic lobes were removed together with the fat bodies, brain membranes and tracheas. Collected samples were rinsed in 0.1M PBS for 10 minutes at room temperature (∼20°C) and then fixed in 4% paraformaldehyde (in 0.1M PBS, pH 7.6) for one hour at room temperature (RT). Brains were washed in PBS 3 times for 15 minutes each at RT, and stored in 0.01% PBS Na-Azide at 4°C. Honey bee brain samples were then prepared with a modified version of the iDISCO protocol (Renier et al., 2014). Samples were dehydrated with increasing concentrations of H2O/methanol (MeOH) (20%, 40%, 60%, 80%, 100%, 100%) for 10 minutes each at RT while gentle shaking. Then samples were incubated for 3 hr at RT, with gentle shaking, in a solution composed by 66% Dichloromethane (DCM) (sigma 270997-250mL, Sigma Aldrich, Italy) and 33% MeOH 100%, and then washed in 100% DCM 5 minutes twice. The optical clearing was performed using 100% DiBenzyl Ether (DBE) (108014-1Kg, Sigma Aldrich, Italy), and incubation was done overnight at RT with gentle shaking.

### TPFM imaging

A custom-made Two-Photon Fluorescence Microscope (TPFM) (Costantini et al., 2021) was used to perform 3D reconstruction of cleared brains. A mode-locked Ti:Sapphire laser (Chameleon, 120 fs pulse width, 80 MHz repetition rate, Coherent, CA) operating at 810 nm was coupled into a custom-made scanning system based on a pair of galvanometric mirrors (LSKGG4/M, Thorlabs, USA). The laser was focused onto the specimen by a 10x objective lens (Plan-Apochromat 10×/0.45, Zeiss, Germany) or a 63x objective (LD C-APOCROMAT 63x/1.15, Zeiss, Germany). The system was equipped with a closed-loop XY stage (U-780 PILine XY Stage System, 135×85 mm travel range, Physik Instrumente, Germany) for radial displacement of the sample and with a closed-loop piezoelectric stage (ND72Z2LAQ PIFOC objective scanning system, 2 mm travel range, Physik Instrumente, Germany) for the displacement of the objective along the z-axis. The fluorescence signal was collected by two independent GaAsP photomultiplier modules (H7422, Hamamatsu Photonics, NJ). Emission filters of (445 ± 40) nm, and (655 ± 40) nm were used to detect the signal, respectively, for tissue autofluorescence and Red Fluorescent MNPs. The instrument was controlled by a custom software, written in LabView (National Instruments, TX) able to acquire a whole sample by performing z-stack imaging of adjacent regions with a voxel size of 0.51×0.51×2 μm. To obtain a single file view of the sample imaged with the TPFM, the acquired stacks were fused together using the ZetaStitcher tool (G. Mazzamuto, “ZetaStitcher: a software tool for high-resolution volumetric stitching” https://github.com/lensbiophotonics/ZetaStitcher) and visualized using Fiji (Schindelin et al., 2012).

### Statistical analysis

Generalised linear mixed models (GLMMs) were used to analyse sucrose responsiveness and the learning and memory performance of control and experimental bees. The GLMMs used a binomial error structure with a logit link function (glmer function of the R package *lme4*, (Bates et al. 2014)). The models were optimised using the iterative “*bobyqa*” or “*nlminbwrap*” algorithm where necessary (Powell, 2009). For the sucrose responsiveness assay, ‘bee response’ was the dependent variable, ‘treatment’ was a fixed factor and ‘sucrose concentration’ was a covariate. For the learning and memory assay, “bee response” was the dependent variable, “treatment” and “CS” were fixed factors, and “conditioning trial” was used as a covariate. Individual identity “ID” was entered as a random factor in all models to account for repeated measures. Interactions were also assessed in all full models. In all cases, the significant model with the highest explanatory power evaluated with the AIC value was selected. To detect differences between the different groups we used Dunnett’s post-hoc tests (*glht* or *lsmeans* function from R package *multcomp*, Bretz et al 2011). Differences in the proportion of bees with specific MTM and eLTM between the experimental and control groups were evaluated using the χ^2^ test. Multiple companions were corrected with Holm-Bonferroni. R 4.0.3 was used for all analyses.

## Results

### Sucrose responsiveness

The graph in Figure 1 illustrates the percentage of PER shown by bees during the six sucrose stimulations when exposed to PMMA, PS, a mixture of the two (MIX), and control groups. The responsiveness of bees in the sucrose assay was found to be dependent on the concentration of sucrose delivered to their antennae. Indeed, as the concentration increased, the proportion of bees exhibiting the PER also increased (GLMM, *Conc*: all P < 0.0001). PMMA alone did not affect sucrose responsiveness compared to controls at any concentration tested (GLMM, treatment: P = 0.96; all three doses, p > 0.05, see Supplementary Table 1). In contrast, overall, the treatment with PS alone reduced sucrose responsiveness (GLMM, P = 0.045). However, post-hoc analysis revealed that only the highest dose tended to reduce it even if not significantly (Dunnett post-hoc test, P = 0.089). Lower doses did not (medium dose: P = 0.68; low dose: P = 0.50, see Supplementary Table 2). The performance of the honey bees exposed to the mixture of the two MPs (MIX) was worse than that of the control bees (the treatment by sucrose concentration interaction was statistically significant at P < 0.0001, see Supplementary Table 3). In this case, the effect was significant for both the high dose (Dunnett post-hoc test, P < 0.0001) and the medium dose (P < 0.0001), but not for the low dose (P = 0.96).

**Figure 1.**
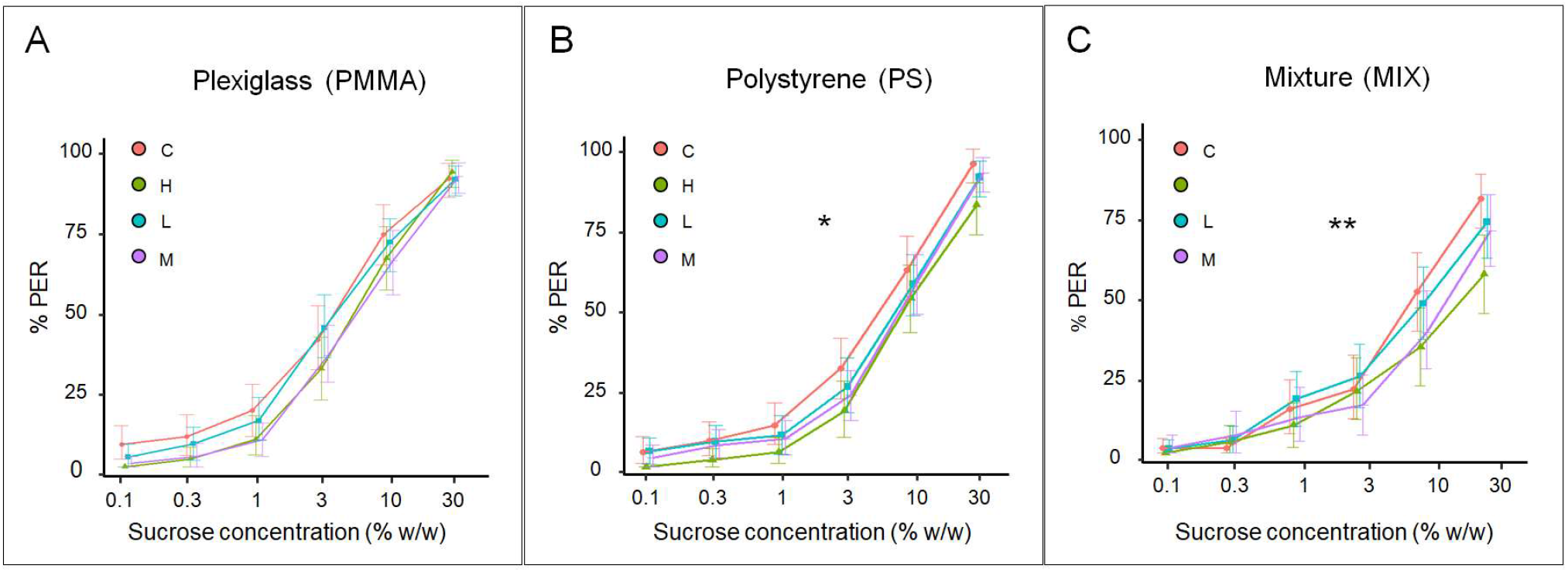
Effect of MPs microparticles on sucrose responsiveness in honey bees. Cumulative percentages of bees exposed to different concentration of MPs (High (H) = green, Low (L) = blue, and Medium (M) = purple line) and control bees (C = red line) showing PER to six sucrose solutions of increasing concentration. (A) Sucrose responsiveness of PMMA-treated bees was not affected compared to the control group, in all concentrations tested. (B) PS treatment affects sucrose responsiveness compared to control as the highest dose tends to reduce it. (C) MIX treatment significantly affects the performance of bees for both the high dose and the medium dose but not for the low dose. (*) *p* < 0.05; (**) *p* < 0.001.

### Associative olfactory learning and memory recall test

The graph in Figure 2 illustrates the percentage of PER shown by bees during five conditioning trials when exposed to PMMA, PS, a mixture of the two (MIX), and control groups. According to the selected GLMM (n°4 for PMMA and PS and GLMM n°6 for MIX, see Supplementary Tables 4, 5, and 6), all groups of bees, increased PER to the odours throughout the conditioning trials (GLMM, *trial*: all P < 0.0001) and, at the same time, showed the ability to discriminate their responses to CS+ and CS-stimuli, regardless of the treatment given (GLMM, *CS*: all P < 0.0001). Learning performance was impaired in bees exposed to PMMA and PS compared to the corresponding controls (GLMM, *treatment*: PMMA: P = 0.02; PS: P = 0.047). The acquisition curves in Figure 2a suggest a dose-dependent effect of PMMA on learning, although post-hoc tests showed no statistically significant differences (Dunnett post-hoc test, PMMA: high dose: P = 0.13; medium dose: P = 0.83; low dose: P = 0.25). PS had a statistically significant effect on the learning phase (Figure 2b) only at the highest dose (Dunnett post-hoc test, GLMM, P = 0.025). There was no significant effect at the lower doses (GLMM, medium dose: P = 0.087; low dose: P = 0.11). Finally, bees exposed to the mixture also showed similar impaired learning compared to controls (Figure 2c), as suggested by the significant treatment by CS interaction (GLMM, P = 0.006). Again, the Dunnett hoc test was not significant (Dunnett post-hoc test, high dose: P = 0.53; medium dose: P = 0.65; low dose: P = 0.86), indicating the absence of an apparent synergistic effect on acquisition performance.

**Figure 2.**
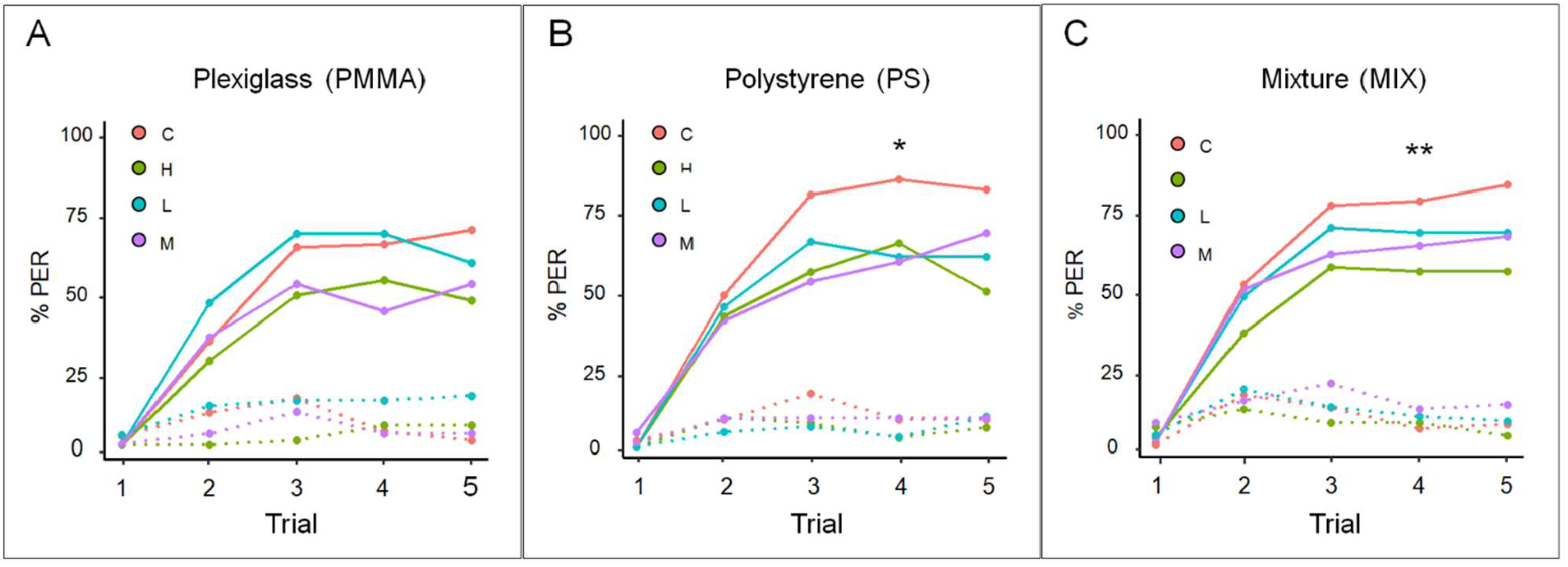
Effect of MPs on associative learning in honey bees. Percentage of PER shown by bees during five conditioning trials when exposed to different concentrations (High (H) = green, Low (L) = blue, and Medium (M) = purple line) of PMMA, PS, a mixture of the two (MIX), and corresponding control groups (C = red line). The solid lines represent the rewarded odorant (CS+) while the dotted lines represent the unrewarded odorant (CS-). (A) Dose-dependent effects of PMMA on bee learning demonstrated in acquisition curves, without statistically significant differences in the post-hoc test. (B) PS treatment causes significant impairment in the learning phase at only the highest dose. (C) Learning phase impairment caused by MIX treatment shown by significant interaction with CS. (*) *p* < 0.05; (**) *p* < 0.01.

Concerning memory retention, overall, the negative effect of PMMA and PS, alone and in combination (MIX), on bee memory retrieval was more pronounced than on memory acquisition (Figure 3). In all cases, both mid-term specific memory (MTM) and early long-term specific memory (eLTM) were impaired compared to controls (χ2 test, MTM: PMMA: X = 9.80, p = 0.02; PS: X = 18.30, p = 0.0004; MIX: X = 13.68, p = 0.003; eLTM: PMMA: X = 12.34, p = 0.006; PS: X = 27.04, p < 0.0001; MIX: X = 18.56, p = 0.0003). In all cases, all doses affected memory recall (C vs High, p < 0.001; C vs Medium, p < 0.014, C vs Low P < 0.02, Figure 3).

**Figure 3:**
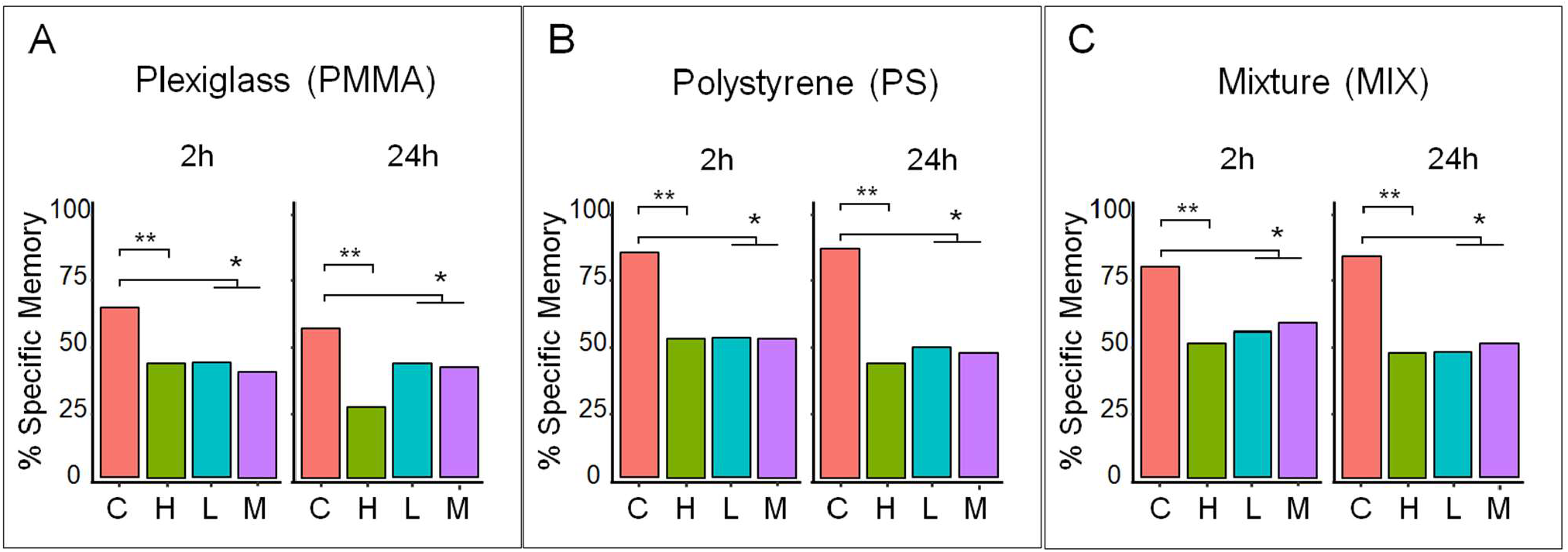
Effect of MPs on mid-term specific memory (MTM) and early long-term specific memory (eLTM) Specific MTM (2h) and eLTM (24h) memory recall of bees exposed to different concentrations of MPs (High (H) = green, Low (L) = blue and Medium (M) = purple bar) and corresponding controls (C = red bar). Exposure to (A) PMMA, (B) PS, and (C) MIX results in significant impairments in both specific memories (MTM and eLTM) compared to controls. All doses of microparticles significantly influence memory recall. (*) *p* < 0.05; (**) *p* < 0.001.

### Brain imaging

Given the small dimension of MPs, the inspection of single thin tissue sections fails to inform on the actual presence of MPs inside the brain. Indeed, during the preparation of the sample, the cutting and mounting processes could modify the location of the MPs, altering a correct evaluation of their location. To finely assess the presence of MPs, we aimed to visualize intact bee brains in 3D with submicron resolution. To this scope we optimized a clearing method to make the brain completely transparent to light modifying the iDISCO technique. The volume was then reconstructed by combining overlapping Z-stacks acquired with Two-Photon Fluorescence Microscopy (TPFM) obtaining the complete 3D reconstruction of the brain with submicron resolution (voxel size = 0,51 × 0,51 × 2 μm) permitting the visualization of the fluorescence MPs in the integrity of the anatomical context. Figure 4 illustrates a 3D reconstruction of a representative dissected honey bee brain before and after the iDISCO clearing acquired with TPFM. Our brain imaging analysis showed that MPs particles, ranging in size from 1 to 5 μm, can reach the foragers’ brains after only three days of oral exposure. The 3D reconstruction performed with TPFM allowed us to clearly distinguish the autofluorescence signal of the brain tissue (in blue) from the red-fluorescing MPs (indicated by white arrows) (Figure 4B). In particular, these microparticles tend to be concentrated mainly in the optic lobes, with aggregation at varying depths within the Medulla tissues (Figure 4C). Our brain analysis focused essentially on a descriptive analysis of this phenomenology and did not aim to quantify the exact number of MPs present in the brain.

**Figure 4:**
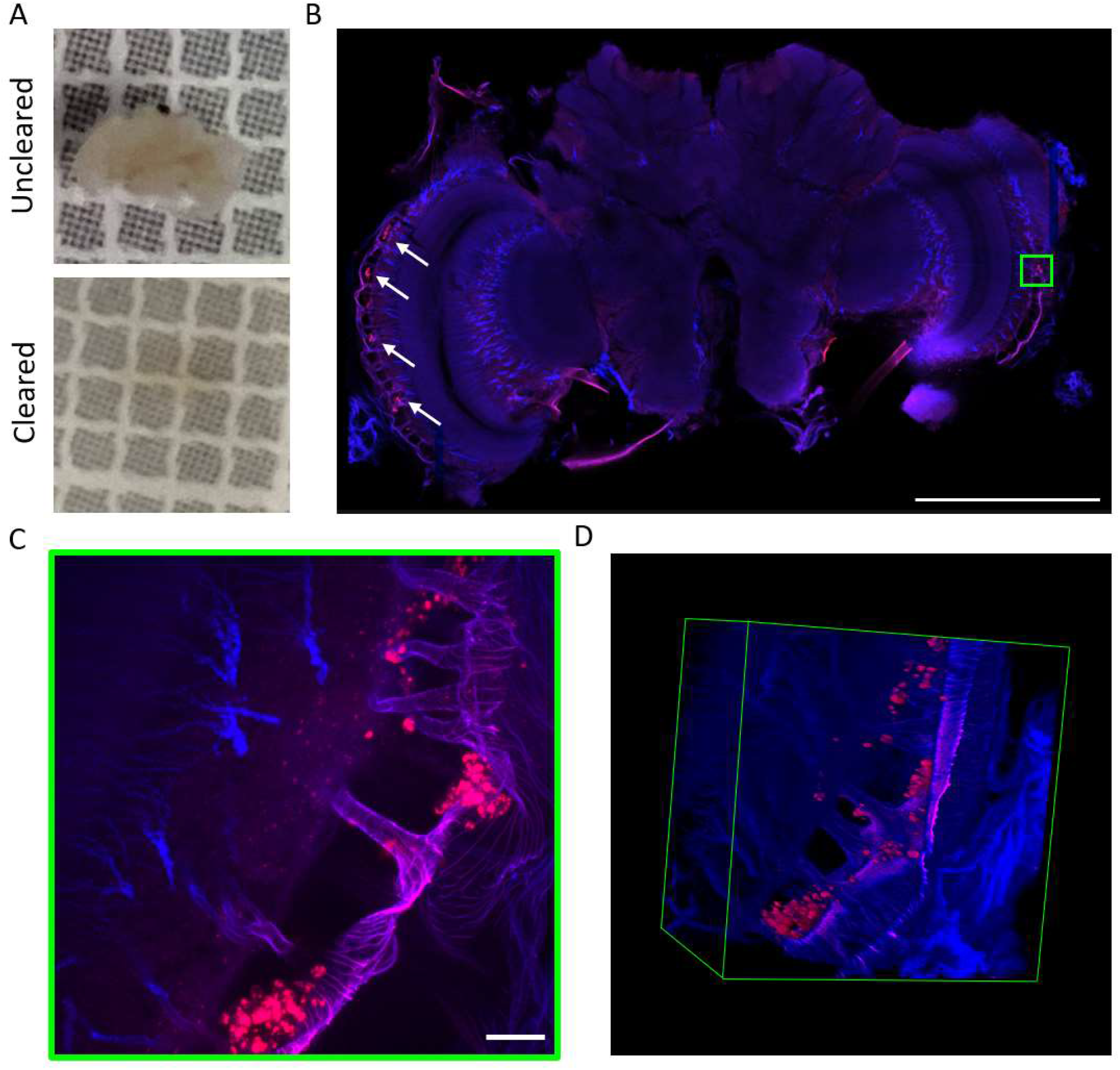
MPs detection in honey bee brain. A) Photos of a dissected brain before and after the iDISCO clearing. B) Single plane (depth ∼200μm) of a 3D reconstruction obtained with TPFM of the whole brain with a 10x objective, resolution of 0,51 × 0,51 × 2 μm^3^. In blue = autofluorescence signal of the tissue; in red = red fluorescence MPs (pointed with white arrows). Scale bar = 1000μm. C) High-resolution inset of the green roi in B acquired with a 63x objective. The image is the maximum intensity projection of a 150 μm depth stack quired with a resolution of 0.17 × 0.17 × 1 μm^3^. Scale bar = 20 μm. D) 3D rendering of the stack in C. Dimensions = 170 × 170 × 150 μm^3^.

## Discussion

Scientific research on the assessment of the effects of MPs in terrestrial systems is still at an early stage but has already partly confirmed that exposure to plastics leads to various negative health consequences on organisms. Recent studies have shown that even important pollinators such as honey bees can enter in contact with MPs in the environment and inadvertently transfer them to other parts of the hive, such as larvae, wax, and honey (Alma et al., 2023). To date, the impact of these polymers on bee behaviour and cognition remains mostly unaddressed with few exceptions (Balzani et al., 2022; Buteler et al., 2022; Liu et al., 2022). To start filling this existing knowledge gap we investigated the impact of MPs on cognitive capacities of *Apis mellifera* and found that and acute oral administration of plexiglass (PMMA), polystyrene (PS), and a combination of both microplastics (MIX) put honey bee foragers at risks. PS reduced sucrose responsiveness, particularly at the highest concentration tested (50 mg L^-1^) which is an essential aspect of honey bee behavior that reflects their ability to detect and respond to nectar sources. PS may therefore potentially affect bee foraging behaviour and foraging efficiency. By contrast, PMMA at any tested dose had no significant effect on sucrose responsiveness. Similar results were found for Polyethylene (PE), which was found to affect only bees’ ability to respond consistently to sucrose but not sucrose responsiveness (Balzani et al., 2022). Unfortunately, PMMA and PS together (MIX) had an even more pronounced negative effect on sucrose reactivity, impairing the bees’ ability to detect and react to sucrose at both high (50 mg L^-1^) and medium (5 mg L^-1^) concentrations. These results could also indicate that the combination of different MP types may have synergistic effects on the cognitive function of *A. mellifera*, which is compromised even at the lowest doses. The negative effects of PS add to other sub-lethal effects on pollinators already reported in the literature for this polymer. These include increased susceptibility to viral infections, damage to digestive system tissues, modifications of genes related to membrane lipid metabolism, detoxification, affect the respiratory system and feeding behaviour (Al Naggar et al., 2023; Deng et al., 2021; C. Wang et al., 2022; K. Wang et al., 2021). Polymer specific effects of PS, PMMA and PE emphasise the significance of considering the distinct characteristics and chemical properties of various polymer types when evaluating their sub-lethal effects. It is plausible to assume, at least with regard to insects, that PS poses a greater risk than other microparticle types (Balzani et al., 2022). Moreover, as also occurs for other stressors such as pesticides and biopesticides or heavy metals (Cappa et al., 2022; Singh et al., 2017), different combination of specific polymers may synergistically amplify the negative of each polymer alone. In addition to the impact on sucrose responsiveness, our study revealed similar concerning outcomes for the other cognitive assessments. Precisely, both PMMA and PS impaired the learning performance of bees when compared to the control group. The impairment in learning was evident during the acquisition phase, as bees exposed to PMMA showed a reduced ability to associate odours with rewards. While the statistical significance of the dose-dependent effect on learning was not established, possibly due to limitations on the power of the post-hoc tests, the data suggest a tendency towards reduced learning ability with higher doses. PS had a significant negative effect on the learning phase, particularly at the highest dose (50 mg/L^-1^). Furthermore, the combination of PMMA and PS also showed impaired learning performance compared to the control group, indicating a cumulative negative effect of both MPs. These findings align with our previous observations on sucrose responsiveness. However, it is important to note that the differences observed in the post-hoc tests did not reach statistical significance. Additionally, PS emerged as the most harmful MPs during the acquisition phase, even when compared to PE microparticles used at the same concentration and under similar experimental conditions and contexts (Balzani et al., 2022). The memory recall tests yielded the most concerning findings, highlighting the detrimental effects of both PMMA and PS, as well as their combination (MIX), on the bee’ cognitive abilities. Our results clearly demonstrated that these MPs adversely affected both medium-term specific memory (MTM) and long-term specific memory (eLTM) when compared to the control group. This suggests that PS, in particular, may impair the ability of bees to retain and retrieve learned information over time. Notably, all tested doses of MPs (50, 5, 0.5 mg L^-1^) had a negative effect on memory recall, highlighting that even ecologically relevant low levels of MPs, commonly encountered in polluted natural environments, can significantly impact cognitive functions in *A. mellifera*.

At present, we have no information regarding the fate and the mechanisms of action of ingested MPs in most organisms. Indeed, a major research gap remains the study of particle translocation to different organs as a function of particle dose and size (Prüst et al., 2020). Our imaging protocol for 3D reconstruction, based on the optimization of a tissue clearing method for bee brains treatment and Two-Photon Fluorescence Microscopy, revealed that MP particles ranging in size from 1-5 μm can penetrate the insect brain within just three days of oral exposure. Specifically, these particles were observed predominantly in the optic lobes, where they aggregated at varying depths within the Medulla tissues. It is important to note that our study primarily focused on providing a descriptive analysis and did not attempt to quantify the number of MPs present in the brain or establish a causal relationship between the observed cognitive impairments and the presence of MPs in the bee brain. However, our findings do demonstrate that MPs can migrate from the gut to the brain neuropils, most likely through the haemolymph, within a short time after exposure. Our results align with the recent scientific evidence that MPs can migrate from the digestive system to different regions of the body. An impressive finding is a study that detected fluorescent MPs in the brain of the velvet crab *Necora puber* after ingestion (Crooks et al., 2019). These particles were found up to 21 days after ingestion, highlighting the persistence and ability of MPs to reach the central nervous system. A similar result was observed in a terrestrial vertebrate species, where 5 μm MPs were found to accumulate in the cortex and hippocampus of aged mice (Liu et al., 2022). The fact that MPs can reach and persist in the central nervous system of bees and other organisms raises the possibility that MP migration may cause mechanical, cellular and biochemical damage in situ. However, there is currently a lack of research and ongoing debate within the scientific community on the potential health risks associated with translocation of MPs (Prüst et al., 2020). Once MPs reach different regions of the insect’s body such as the gut, haemolymph, Malpighian tubules, or fat bodies in the abdominal cavity, it is plausible that they undergo metabolic degradation or detoxification through enzymatic processes or interactions with the insect’s gut microbiome. These metabolic processes could potentially lead to the production of secondary metabolites that induce oxidative stress, cellular damage and an increased susceptibility to neurological disorders. In addition, the activation of the immune system by these MPs or their secondary metabolites may directly affect cognitive abilities (Mallon et al., 2003; Riddell & Mallon, 2006). Consequently, the observed behavioural changes in our bees induced by MPs may also be due to activation of the immune system at the peripheral level rather than occurring directly in the brain. Nevertheless, although limited, existing evidence in mammals suggests that exposure to MPs may directly interfere with the activity of acetylcholinesterase and disrupt the balance of neuromodulators and neurotransmitters (Prüst et al., 2020). To gain further insights into the underlying mechanisms of action of MPs on cognition, future studies in bees should focus on addressing this aspect.

In conclusion, our findings are alarming and raise concerns about the potential long-term consequences of exposure to MPs on honey bees, as well as other beneficial insects or arthropods. The research conducted in our study focused mainly on the effects of acute exposure; however, there is a need for a deeper understanding of the long-term effects of chronic exposure to MPs, particularly polystyrene (PS), which is known to be the major contributor to a number of sub-lethal effects in bees. A deeper understanding of the effects can be achieved through additional studies that investigate the accumulation and persistence of MPs in bees’ brains and how this affects their sensory systems, behavioural patterns, and cognitive mechanisms.

## Author Contributions

D.B. conceptualized and designed the study, E.P., L.P. and F.F. performed behavioural experiments. I.C. optimized the clearing method, performed the TPFM imaging and analysed the data. D.B. performed statistical analysis. F.S.P. and D.B. provided funding and equipment for imaging analysis. E.P. wrote the first draft of the manuscript. D.B., E.P. and I.C. contributed to writing the final version of the manuscript.

