## Supplementary Tables for "Microplastics reach the brain and interfere with honey bee cognition"

**Table S1:** Results of Generalized Linear Mixed Models for the sucrose responsiveness assay (PS)

| **SRS - PS** | Dependent Variable | | | |
|---|---|---|---|---|
| | Response | | | |
| | (1) | (2) | (3) | (4) |
| Treatment High | | -0.446***<br>(0.164) | -1.741**<br>(0.827) | -3.232<br>(2.187) |
| Treatment Low | | -0.134<br>(0.155) | -0.541<br>(0.793) | 0.219<br>(1.815) |
| Treatment Medium | | -0.179<br>(0.154) | -0.714<br>(0.788) | -1.509<br>(1.898) |
| Sucrose Concentration | | | 3.078***<br>(0.139) | 3.016***<br>(0.276) |
| Treatment High: Sucrose Concentration | | | | 0.315<br>(0.436) |
| Treatment Low: Sucrose concentration | | | | -0.167<br>(0.370) |
| Treatment Medium :Sucrose Concentration | | | | 0.177<br>(0.388) |
| Constant | -0.837***<br>(0.057) | -0.650***<br>(0.114) | -13.509***<br>(0.840) | -13.239***<br>(1.334) |
| Akaike Inf. Crit. | 2,776.448 | 2,774.730 | 1,340.619 | 1,343.411 |
| Note | *p<0.1; **p<0.05; ***p<0.01 | | | |

**Table S2:** Results of Generalized Linear Mixed Models for the sucrose responsiveness assay (PMMA)

| **SRS - PMMA** | Dependent Variable | | | |
|---|---|---|---|---|
| | Response | | | |
| | (1) | (2) | (3) | (4) |
| Treatment High | | -0.280* (0.146) | -0.420 (1.169) | -2.662 (6.794) |
| Treatment Low | | -0.059 (0.139) | 0.127 (1.227) | -3.123 (6.976) |
| Treatment Medium | | -0.253* (0.141) | -0.348 (1.161) | -9.762 (6.833) |
| Sucrose Concentration | | | 15.227*** (0.951) | 14.331*** (1.411) |
| Treatment High: Sucrose Concentration | | | | 0.460 (1.428) |
| Treatment Low: Sucrose concentration | | | | 0.692 (1.440) |
| Treatment Medium :Sucrose Concentration | | | | 2.051 (1.443) |
| Constant | -0.541*** (0.051) | -0.395*** (0.101) | -68.109*** (4.464) | -63.904*** (6.781) |
| Akaike Inf. Crit. | 2,755.069 | 2,755.366 | 1,072.253 | 1,075.946 |
| Note | *p<0.1; **p<0.05; ***p<0.01 | | | |

**Table S3:** Results of Generalized Linear Mixed Models for the sucrose responsiveness assay (MIX)

| **SRS - MIX** | Dependent Variable | | | |
|---|---|---|---|---|
| | Response | | | |
| | (1) | (2) | (3) | (4) |
| Treatment High | | -0.541**<br>(0.269) | -21.458***<br>(1.816) | 5.202<br>(9.053) |
| Treatment Low | | -0.029<br>(0.245) | -0.175<br>(1.285) | 6.345<br>(10.128) |
| Treatment Medium | | -0.284<br>(0.268) | -0.341<br>(1.298) | -8.151<br>(10.426) |
| Sucrose Concentration | | | 19.428***<br>(1.122) | 21.100***<br>(1.913) |
| Treatment High: Sucrose Concentration | | | | -7.955***<br>(1.852) |
| Treatment Low: Sucrose concentration | | | | -1.424<br>(2.219) |
| Treatment Medium :Sucrose Concentration | | | | -2.270<br>(2.077) |
| Constant | -1.289***<br>(0.101) | -1.098***<br>(0.178) | -106.476***<br>(6.296) | -95.361***<br>(8.743) |
| Akaike Inf. Crit. | 1,588.116 | 1,588.785 | 680.031 | 673.377 |
| Note | *p<0.1; **p<0.05; ***p<0.01 | | | |

**Table S4:** Results of Generalized Linear Mixed Models for the learning assay (PS)

| **Learning - PS** | Dependent Variable | | | | | |
|---|---|---|---|---|---|---|
| | RespAQ | | | | | |
| | (1) | (2) | (3) | (4) | (5) | (6) |
| Group H | | -0.614*** (0.236) | -0.682*** (0.261) | -0.998*** (0.383) | -0.192 (0.561) | -0.129 (0.564) |
| Group L | | -0.497** (0.236) | -0.554** (0.262) | -0.814** (0.384) | -0.339 (0.564) | -0.288 (0.572) |
| Group M | | -0.470** (0.234) | -0.523** (0.260) | -0.769** (0.381) | -0.433 (0.563) | -0.359 (0.566) |
| Sequence | | | 0.501*** (0.037) | 0.729*** (0.050) | 0.850*** (0.091) | 0.922*** (0.102) |
| CSB | | | | -3.067*** (0.156) | -3.078*** (0.157) | -3.578*** (0.309) |
| Group H: Sequence | | | | | -0.244** (0.125) | -0.339** (0.135) |
| Group L: Sequence | | | | | -0.146 (0.125) | -0.206 (0.139) |
| Group M: Sequence | | | | | -0.104 (0.125) | -0.208 (0.137) |
| Group H: CSB | | | | | | 0.772* (0.412) |
| Group L :CSB | | | | | | 0.397 (0.423) |
| Group M :CSB | | | | | | 0.816** (0.408) |
| Constant | -1.181*** (0.088) | -0.780*** (0.166) | -2.372*** (0.222) | -1.929*** (0.309) | -2.320*** (0.395) | -2.379*** (0.406) |
| Akaike Inf. Crit. | 2,979.859 | 2,978.016 | 2,781.495 | 2,179.937 | 2,181.971 | 2,182.739 |
| Note | *p<0.1; **p<0.05; ***p<0.01 | | | | | |

**Table S5:** Results of Generalized Linear Mixed Models for the learning assay (PMMA)

| **Learning - PMMA** | Dependent Variable | | | | | |
|---|---|---|---|---|---|---|
| | RespAQ | | | | | |
| | (1) | (2) | (3) | (4) | (5) | (6) |
| Group H | | -0.507* (0.259) | -0.564** (0.285) | -0.762** (0.387) | -0.794 (0.594) | -0.801 (0.599) |
| Group L | | 0.176 (0.250) | 0.187 (0.276) | 0.261 (0.376) | 0.301 (0.555) | 0.310 (0.554) |
| Group M | | -0.430 (0.264) | -0.478 (0.291) | -0.636 (0.394) | -0.286 (0.595) | -0.277 (0.597) |
| Sequence | | | 0.503*** (0.040) | 0.687*** (0.050) | 0.710*** (0.089) | 0.726*** (0.093) |
| CSB | | | | -2.859*** (0.159) | -2.862*** (0.159) | -3.018*** (0.287) |
| Group H: Sequence | | | | | 0.008 (0.131) | 0.008 (0.139) |
| Group L: Sequence | | | | | -0.012 (0.122) | -0.056 (0.127) |
| Group M: Sequence | | | | | -0.102 (0.131) | -0.114 (0.137) |
| Group H: CSB | | | | | | -0.019 (0.435) |
| Group L :CSB | | | | | | 0.457 (0.383) |
| Group M :CSB | | | | | | 0.103 (0.436) |
| Constant | -1.569*** (0.102) | -1.385*** (0.179) | -3.037*** (0.244) | -2.752*** (0.316) | -2.829*** (0.401) | -2.838*** (0.404) |
| Akaike Inf. Crit. | 2,772.640 | 2,769.139 | 2,594.282 | 2,107.861 | 2,113.044 | 2,117.128 |
| Note | *p<0.1; **p<0.05; ***p<0.01 | | | | | |

**Table S6:** Results of Generalized Linear Mixed Models for the learning assay (MIX)

| **Learning - MIX** | Dependent Variable | | | | | |
|---|---|---|---|---|---|---|
| | RespAQ | | | | | |
| | (1) | (2) | (3) | (4) | (5) | (6) |
| Group H | | -0.600**<br>(0.238) | -0.660**<br>(0.261) | -0.936**<br>(0.371) | -0.243<br>(0.531) | -0.176<br>(0.536) |
| Group L | | -0.195<br>(0.236) | -0.217<br>(0.260) | -0.319<br>(0.368) | 0.050<br>(0.522) | 0.125<br>(0.529) |
| Group M | | -0.203<br>(0.235) | -0.226<br>(0.258) | -0.328<br>(0.365) | -0.055<br>(0.522) | 0.003<br>(0.526) |
| Sequence | | | 0.479***<br>(0.035) | 0.676***<br>(0.045) | 0.772***<br>(0.080) | 0.872***<br>(0.092) |
| CSB | | | | -2.908***<br>(0.141) | -2.916***<br>(0.142) | -3.699***<br>(0.287) |
| Group H: Sequence | | | | | -0.209*<br>(0.115) | -0.330***<br>(0.125) |
| Group L: Sequence | | | | | -0.113<br>(0.113) | -0.222*<br>(0.125) |
| Group M: Sequence | | | | | -0.084<br>(0.114) | -0.227*<br>(0.123) |
| Group H: CSB | | | | | | 1.051***<br>(0.387) |
| Group L :CSB | | | | | | 1.051***<br>(0.387) |
| Group M :CSB | | | | | | 1.238***<br>(0.371) |
| Constant | -1.141***<br>(0.088) | -0.896***<br>(0.162) | -2.422***<br>(0.213) | -1.970***<br>(0.288) | -2.279***<br>(0.360) | -2.352***<br>(0.373) |
| Akaike Inf. Crit. | 3,390.390 | 3,389.883 | 3,186.291 | 2,550.122 | 2,552.776 | 2,545.733 |
| Note | *p<0.1; **p<0.05; ***p<0.01 | | | | | |
